# Determinants of Cancer-specific Quality of Life in Veteran Lung Cancer Survivors Eligible for Long-Term Cure

**DOI:** 10.1101/518910

**Authors:** Duc Ha, Andrew L. Ries, Jeffrey J. Swigris

## Abstract

**Rationale/Objective:** Quality of life (QoL) is an important issue in lung cancer survivors. We aimed to identify determinants of QoL in lung cancer survivors eligible for long-term cure.

**Methods:** We performed an exploratory analysis of a cross-sectional study of consecutive lung cancer survivors who completed curative-intent treatment ≥1 month previously. Variables tested included demographic, clinical, physiologic, and symptom-specific patient-reported outcome measures. We defined the primary outcome as a previously-validated cancer-specific QoL measure – the European Organization for Research and Treatment of Cancer QoL Questionnaire Core 30 (C30) summary score. We also verified our findings with the C30 global health status/QoL subscale and a summated score of lung cancer-specific QoL from the EORTC-Lung Cancer Module 13.

**Results:** In 75 enrolled participants, measures of fatigue, depression, sleep difficulties, and dyspnea were statistically significant determinants of the C30 summary score in multivariable linear regression analyses. Together, these four symptoms accounted for approximately 85% of the variance in cancer-specific QoL (p<0.001). When we verified our findings with global QoL and lung cancer-specific QoL, fatigue and dyspnea were consistent determinants of QoL.

**Conclusions:** We found four symptoms – dyspnea, fatigue, depression, and sleep difficulties – that are important determinants of and together accounted for almost all of the variance in cancer-specific QoL in lung cancer survivors eligible for long-term cure. These findings have implications to reduce symptom burden and improve function and QoL in these patients.

## Introduction

Lung cancer is the second-most commonly diagnosed cancer in the United States (US).^1^ Up to 50% of cases of non-small cell lung cancer (NSCLC) are diagnosed at local or locally-advanced stage,^2^ the standard treatment of which uses a combination of surgical resection, radio-ablation therapy, and chemotherapy aimed at achieving a cure.^3–5^ As of 2016, there were >500,000 lung cancer survivors in the US.^6^ The number of lung cancer survivors eligible for long-term cure is increasing due to a potential stage shift^7^ from implementation of lung cancer screening in high-risk individuals^8^ and adoption and implementation of screening programs across healthcare systems in the US.

Threats to quality of life (QoL) among lung cancer survivors include physical and emotional effects of aging, health behaviors, comorbid conditions, and lung cancer and its treatment. The median age at lung cancer diagnosis is 70 years, compared to 62 for breast, 66 for prostate, and 67 for colorectal cancers.^9^ Approximately 30% of lung cancer survivors continue to smoke^10^ and 80% do not meet physical activity recommendations.^11^ The most common comorbidities in patients with lung cancer, all of which can progressively and negatively impact health, include chronic obstructive pulmonary disease (COPD, in 53% of patients), diabetes (16%), and congestive heart failure (HF, 13%).^12^ Each therapeutic modality (surgery, radio- and chemotherapy) carries risk of adverse effects: in patients undergoing lung cancer resection surgery, these include perioperative pulmonary^13^ and/or cardiac^14^ complications and surgery-related loss of lung function.^15^ After radio-ablation, some degree of focal pulmonary fibrosis will occur; 5-15% of patients experience acute pneumonitis, and a significant minority are at risk for progressive pulmonary fibrosis and respiratory failure.^16,17^ The potential acute and long-term effects of chemotherapy are numerous^18^ and, like surgery and radiotherapy, pose threats to physical functioning, emotional well-being, independence and social interactions in lung cancer survivors.

Quality of life is an important outcome to lung cancer survivors and more important than length of survival for many.^19^ Identifying determinants of QoL in this patient population may reveal factors that can be mitigated. In this project, we aimed to identify determinants of QoL in lung cancer survivors eligible for long-term cure.

## Methods

### Study Overview

We previously performed an Institutional Review Board-approved, cross-sectional study to assess functional capacity and patient-reported outcomes (PROs) in lung cancer survivors following curative-intent therapy.^20^ In the current study, we enrolled additional participants to identify determinants of QoL. Between August 2016 and April 2018, we recruited patients by mailing informational letters to consecutive patients whose lung cancer was diagnosed and/or managed at the VA San Diego Healthcare System (VASDHS) since October 2010. We followed up with a telephone call one week later to gauge their interest, obtained written informed consent from all participants, and followed guideline recommendations to report our findings.^21^

### Participants

We enrolled a convenience sample of lung cancer survivors who completed curative-intent treatment, defined as lung cancer resection surgery, definitive radio-ablation, or chemoradiation for stage I-IIIA disease ≥1 month previously. We excluded patients with severe dementia (n=2), bilateral below-knee amputation (n=2), quadriplegia (n=1), or active systemic treatment for other cancers (n=4). Following exclusion, we enrolled 87% of all eligible patients.^22^

### Variables

We abstracted demographic, clinical, lung cancer-specific, and physiologic data from the medical record. We assessed patient-reported symptoms, including fatigue, anxiety/depression, sleep difficulties, dyspnea, and pain by using the Brief Fatigue Inventory (BFI),^23^ Hospital Anxiety and Depression Scale (HADS),^24^ Pittsburgh Sleep Quality Index (PSQI),^25^ University California San Diego Shortness of Breath Questionnaire (SOBQ),^26^ and European Organization for Research and Treatment of Cancer QoL Questionnaire Lung Cancer Module 13 (LC13),^27^ respectively. All questionnaire assessments were obtained at the same time, on printed forms without any modifications.

### Primary and Secondary Outcomes

The primary outcome, cancer-specific QoL, was defined as the EORTC-QLQ-Core 30 (C30) summary score. The C30 consists of 30 items on 15 different domains of health specific to the cancer patient population.^28^ The summary score is calculated as the mean of 13 of the 15 scales: physical function, role function, social function, emotional function, cognitive function, fatigue, nausea/vomiting, dyspnea, sleep disturbances, appetite loss, constipation, and diarrhea (the global QoL and financial difficulty scales are excluded) and has been previously used in the lung cancer population.^22^ C30 summary scores range from 0 to 100, and higher scores connote better QoL. We used the global health status/QoL subscale of the C30^28^ and lung cancer-specific QoL from the LC13, scored as a summation of responses as reported by previous literature^29,30^ to verify our findings. LC13 scores range from 12 to 48, and higher scores correspond to worse QoL.

### Statistical Analyses

We summarized descriptive statistics as appropriate. We used linear regression analyses to identify determinants of the outcomes. In multivariable models, we used stepwise, backwards selection modeling starting with all variables with p <0.05 in univariate analyses, and additionally forced age,^31,32^ sex,^31,32^ race,^19^ and smoking status^31,32^ into the model for QoL regardless of statistical significance. We defined statistical significance as p <0.05 in two-tailed tests. All data were managed using REDCap electronic data capture tools hosted at the UCSD Clinical and Translational Research Institute^33^ and analyzed using IBM^®^ SPSS^®^ Statistics software version 25.

## Results

### Participants

We enrolled 75 lung cancer survivors whose baseline characteristics are as described in Table 1. Typical for a VA population, most were white males with a history of tobacco use, concomitant COPD, and had undergone either surgical resection or definitive radio-ablation for treatment of stage I NSCLC. The median time since treatment completion was 12 months.

### Physiological and Patient-Reported Outcome Assessments

Results of physiological and patient-reported outcome (PRO) assessments are as summarized in Table 2. The most common abnormal symptoms^34^ were sleep difficulties, dyspnea, and pain. The mean (SD) LC13 score was 75.5 (17.5),^35^ suggesting significant impairment in cancer-specific QoL.^36^

### Determinants of Cancer-specific QoL (C30 summary score; primary outcome variable)

Determinants of C30 summary score are bolded in Table 3A, with selected variables displayed in Figure 1A-C. In models adjusting for age, sex, race, and smoking history, we identified fatigue, depression, sleep quality, and dyspnea to be independent predictors of cancer-specific QoL. In all models, these four symptoms explained approximately 85% of the variance (p <0.001) in C30 summary score (Table 3B, Models 1-3). Fatigue and dyspnea were associated with <u>global</u> QoL and <u>lung cancer-specific</u> QoL (E-Tables 1, 2A, 2B), and anxiety with lung cancer-specific QoL (Table 2B).^37^

## Discussion

In this cross-sectional study of consecutive veteran lung cancer survivors eligible for long-term cure, we found symptoms of fatigue, depression, sleep difficulties, and dyspnea to be significant determinants of cancer-specific QoL that, together, accounted for approximately 85% of its variance. We also verified fatigue and dyspnea as determinants of global and lung cancer-specific QoL. To the best of our knowledge, this study is the first to examine the relationship between symptom-specific PROs and a validated cancer-specific QoL summary score in this patient population. The results of our study have implications in lung cancer survivorship.

A cancer survivor is defined by the National Coalition for Cancer Survivorship and American Cancer Society as a person diagnosed with cancer starting from the time of diagnosis to end of life.^38,39^ In a seminal report on issues related to cancer survivorship, the Institute of Medicine emphasized care in the post-treatment phase to include care planning, supportive services, and management and prevention of late treatment effects.^40^ QoL is an important outcome for lung cancer survivors following curative-intent treatment^41^ and, among other factors, may be affected by the physical and psychosocial consequences of lung cancer and its treatment.

The mean C30 summary score of 75.5 in our study is similar to another cohort of 66 lung cancer patients undergoing anatomic lung cancer resection surgery.^36^ Our study adds to existing literature by identifying four important symptoms which account for a substantial amount of the variance in cancer-specific QoL. These results reveal potential opportunities for impactful clinical management strategies: initiating and/or optimizing therapies that target fatigue, dyspnea, poor sleep quality and mood disturbance could substantially improve QoL in these patients.

Our results are also similar to those from another study^42^ designed to assess the impact of symptom burden on generic QoL in 183 post-surgical NSCLC survivors. In that study, investigators found that pain, fatigue, dyspnea, and depression/anxiety were associated with physical QoL and having three or more symptoms was associated with mental QoL.^42^ When we included LC13 subscales in our analyses (results not shown), we also found bodily pain, cough, hemoptysis, sore mouth, and dysphagia to be associated with cancer-specific QoL; however in multivariable models, only sore mouth remained an independent determinant of cancer-specific QoL. Similarly, in another study of 447 post-surgical lung cancer survivors, fatigue and pain were associated with poor overall QoL at short- and long-term (≥5 years) follow-up.^43^

Along with the Functional Assessment of Cancer Therapy–General^44^/Lung^45^ (FACT-G/L), the C30^28^/LC13^27^ questionnaires are the most commonly used cancer/lung cancer-specific PRO instruments. Despite their common use, fatigue and dyspnea, two of the most common symptoms in lung cancer survivors,^20,37,46,47^ are assessed by only one and three items, respectively, on the C30, and only three and zero questions, respectively, on the LC13. Supported by previous literature, we calculated a summated lung cancer-specific QoL score from the LC13;^29,30^ however, this score only includes some of the symptoms associated with lung cancer and does not include scales on function, emotional and psychological well-being, and other symptoms known to be important for lung cancer-related health (e.g. fatigue). While there are ongoing efforts to create a more comprehensive lung cancer-specific instrument with a summary score^48^, it is not currently available. Thus, we used existing, symptom-specific questionnaires to incorporate these two important symptoms.

In contrast to other studies, in our cohort, age,^31,32^ sex,^31,32^ race,^19^ and smoking status^31,32^ were not significantly associated with cancer-specific QoL; this may be due to a lack of statistical power and/or inclusion of symptom-specific PROs in our models which may indirectly adjust for these variables.

Compared to usual care, physical exercise has been shown to be effective in improving physical function and QoL in cancer survivors.^49^ However, out of the 34 randomized clinical trials included in a recent systematic review, only one (N=51) focused on lung cancer survivors.^49^ Thus, although there is rationale for encouraging physical exercise in lung cancer survivors, its beneficial effects on cancer-specific, lung cancer-specific and global QoL requires additional investigation.

Lung cancer survivors are somewhat different from other cancer populations due to a higher median age at diagnosis, lifetime exposure to tobacco, and prevalence of pulmonary and cardiovascular comorbidities. In addition, the effects of curative-intent lung cancer treatment have unique effects on cardiopulmonary health since part of the therapy requires local destruction and/or removal of lung tissue which may impede cardiopulmonary function, exercise capacity, and dyspnea symptoms. As such, compared to a propensity-matched sample of 1,000 individuals from a general population in Korea,^50^ dyspnea scores were higher in 830 disease-free, post-surgical lung cancer survivors. Adjusted dyspnea scores were also higher in those treated with combined-modality therapy compared to those treated by surgery alone, suggesting treatment-related effects.^50^ Equally important, those with cardiac or pulmonary comorbidity had a higher dyspnea score than those without any comorbidity, suggesting a cardiopulmonary comorbidity effect.^50^ Together, these data suggest that to improve exercise, function and QoL in lung cancer survivors, medical therapy for cardiopulmonary disease and other comorbid conditions may need to be optimized to alleviate symptom burden before patients can participate in regular exercise to improve function and QoL.

Our study has limitations. First, we did not include other determinants of health which may impact cancer-specific QoL, including marital and socio-economic status, living condition, and education level. Second, we did not have a comparison group; therefore, we cannot make inferences on differences in determinants of cancer-specific QoL in lung cancer compared to other cancer survivors. Third, the descriptive, cross-sectional, and observational nature of our study provides no insight into the underlying physiobiological mechanisms or the effects of lung cancer and its treatment on QoL. Fourth, our findings may have limited generalizability, due to it being a single-institutional study involving a predominantly white male veteran patient population with significant tobacco exposure and higher prevalence of COPD and psychiatric illness.

The strengths of our study include a detailed list of comorbidities and physiological measures of cardiopulmonary health. In addition, all PRO assessments were performed in-person by one observer (DH), maximizing the completeness and accuracy of the data collected and minimizing inter-observer variability. Also, we verified our findings using two other QoL measures, strengthening the validity of our conclusions. Last, we identified determinants which account for almost all of cancer-specific QoL, facilitating interpretation and translation into the clinical setting.

We conclude that in this cross-sectional study of consecutive lung cancer survivors eligible for long-term cure, we identified several important determinants of cancer-specific QoL which may have clinical implications to improve lung cancer survivorship. Efforts to improve QoL in these patients may need to focus on reducing fatigue, dyspnea, and possibly anxiety/depression symptoms and/or sleep difficulties to be effective.

**Table 1:**
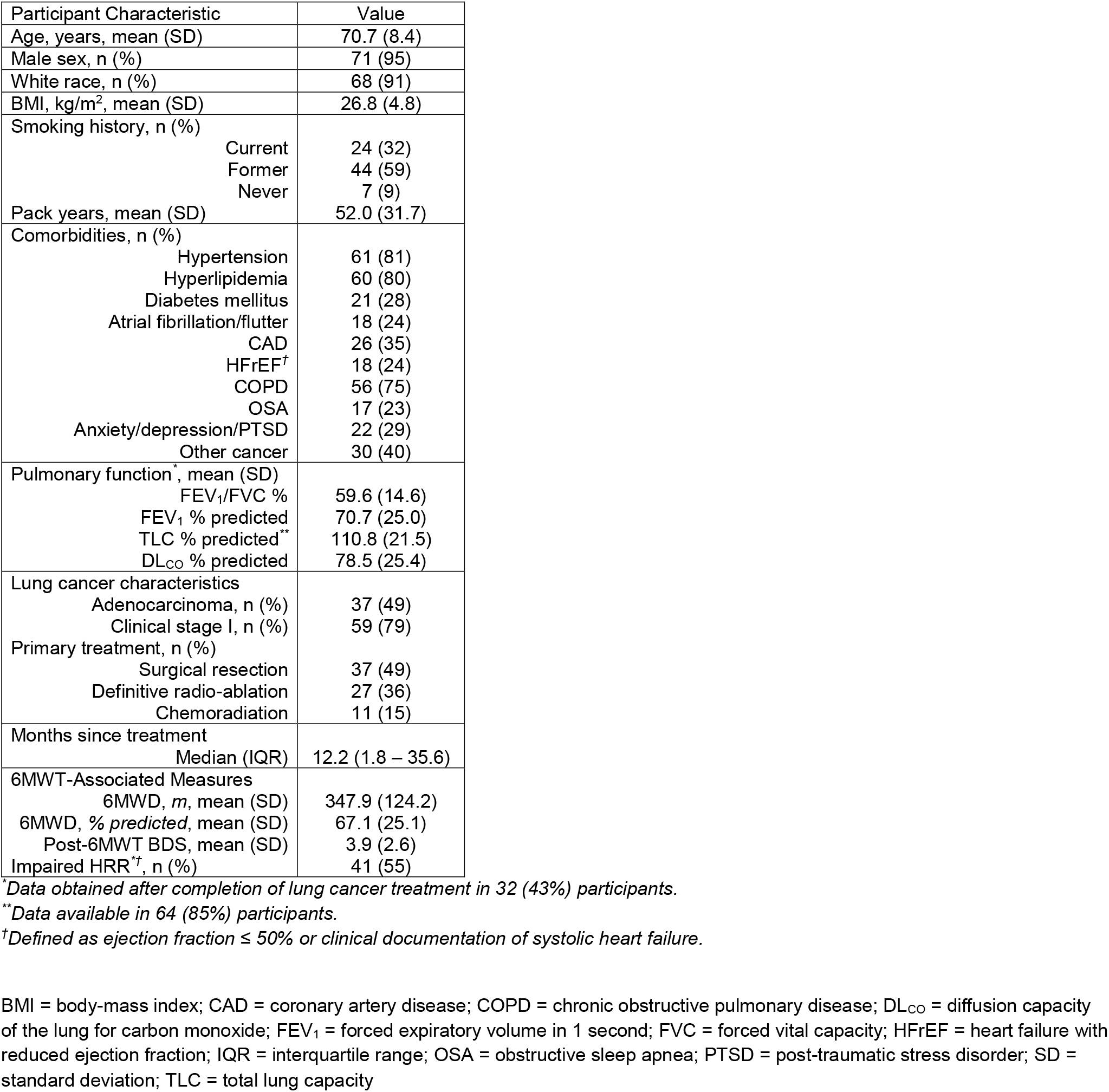
Participant Characteristics

**Table 2:** Patient-Reported Outcome Assessments

| Patient-Reported Outcome Assessments |  |  |
| --- | --- | --- |
| Symptom-specific PRO, mean (SD) |  |  |
| BFI |  | 25.7 (23.3) |
| HADS – Anxiety |  | 4.6 (4.4) |
| HADS – Depression |  | 5.8 (4.4) |
| PSQI |  | 8.2 (4.7) |
| SOBQ |  | 34.8 (25.6) |
| LC13 |  |  |
| Dyspnea |  | 35.3 (26.2) |
| Coughing |  | 39.6 (26.7) |
| Hemoptysis |  | 4.4 (16.7) |
| Sore mouth |  | 3.1 (11.2) |
| Dysphagia* |  | 9.5 (21.0) |
| Peripheral neuropathy |  | 17.8 (27.6) |
| Alopecia |  | 5.3 (13.5) |
| Pain in chest |  | 16.4 (25.3) |
| Pain in arm or shoulder |  | 25.3 (29.9) |
| Pain in other parts* |  | 34.7 (33.3) |
| Quality of Life<sup>‡</sup> scores, mean (SD) |  |  |
| C30 summary score<sup>35</sup> |  | 75.5 (17.5) |
| Global QoL<sup>28</sup> |  | 62.7 (23.3) |
| Lung cancer-specific QoL<sup>29,30</sup> |  | 19.7 (4.7) |

\*Data available in 74 participants for HRR, dysphagia, and 72 participants for pain in other parts.
<sup>†</sup>Defined as HRR ≤ 12 beats at 1-minute following 6MWT.<sup>51,52</sup>
<sup>‡</sup>Defined as C30 summary score for cancer-specific QoL (range 0 – 100);<sup>35</sup> C30 global health status/QoL for global QoL (range 0 – 100);<sup>28</sup> and summated LC13 score from questions 31 – 42 for lung cancer-specific QoL (maximum score = 48, indicating worst possible QoL).<sup>29,30</sup>
6MWD = six-minute walk distance; 6MWT = six-minute walk test; BDS = Borg Dyspnea Score; BFI = Brief Fatigue Inventory; C30/LC13 = European Organization for Research and Treatment of Cancer QoL Core 30/Lung Cancer Module 13; HADS = Hospital
Anxiety and Depression Scale; HRR = heart rate recovery; PRO = patient-reported outcome; PSQI = Pittsburg Sleep Quality Index; QoL = quality of life; SD = standard deviation; SOBQ = University of California San Diego Shortness of Breath Questionnaire

**Table 3A:** Predictors of cancer-specific QoL (C30 summary score)

| Variable | $\beta$ | R<sup>2</sup> | P value |
| --- | --- | --- | --- |
| Age, year | 0.44 | 0.05 | 0.07 |
| Sex (F/M) | -6.08 | 0.01 | 0.50 |
| White race (N/Y) | -8.01 | 0.02 | 0.25 |
| BMI, kg/m<sup>2</sup> | -0.07 | 0.00 | 0.88 |
| Smoking history | F-statistics | 0.05 | 0.13 |
| Comorbidities (N/Y) |  |  |  |
| Diabetes mellitus | 2.98 | 0.01 | 0.51 |
| Atrial fibrillation/flutter | 2.61 | 0.004 | 0.59 |
| CAD | 0.06 | 0.00 | 0.99 |
| **HFrEF** | **9.40** | **0.05** | **0.04** |
| COPD | -1.60 | 0.002 | 0.73 |
| **OSA** | **10.8** | **0.07** | **0.03** |
| **Anxiety/depression/PTSD** | **16.2** | **0.18** | **<0.001** |
| Other cancer | -3.08 | 0.01 | 0.46 |
| Pulmonary function |  |  |  |
| FEV<sub>1</sub> % predicted, each | 0.08 | 0.01 | 0.34 |
| TLC % predicted, each | -0.10 | 0.02 | 0.32 |
| DL<sub>CO</sub> % predicted, each | 0.13 | 0.03 | 0.12 |
| Lung cancer characteristics |  |  |  |
| Clinical stage I (N/Y) | 0.73 | 0.00 | 0.89 |
| Primary treatment | F-statistics | 0.04 | 0.23 |
| Months since treatment, each | 0.04 | 0.01 | 0.48 |
| 6MWT-associated measures |  |  |  |
| **6MWD, each m** | **0.05** | **0.11** | **0.01** |
| **Post-6MWT BDS, each point** | **-3.81** | **0.33** | **<0.001** |
| HR change, each beats/min | 0.16 | 0.01 | 0.34 |
| O<sub>2</sub> saturation change, each % | 0.04 | 0.00 | 0.92 |
| HRR, each beat | -0.16 | 0.004 | 0.61 |
| Symptom-specific PRO, each point |  |  |  |
| **BFI** | **-0.65** | **0.74** | **<0.001** |
| **HADS anxiety** | **-2.51** | **0.39** | **<0.001** |
| **HADS depression** | **-3.20** | **0.64** | **<0.001** |
| **PSQI** | **-2.32** | **0.35** | **<0.001** |
| **SOBQ** | **-0.53** | **0.60** | **<0.001** |

**Bolded** variables indicate statistically significant associations at $p < 0.05$ and selected to enter MVAs.
6MWD = six-minute walk distance; 6MWT = six-minute walk test; BDS = Borg Dyspnea Score; BFI = Brief Fatigue Inventory; C30 = European Organization for Research and Treatment of Cancer QoL Core 30; CAD = coronary artery disease; COPD = chronic obstructive pulmonary disease; $DL_{CO}$ = diffusion capacity of the lung for carbon monoxide; $FEV_1$ = forced expiratory volume in 1 second; HADS = Hospital Anxiety and Depression Scale; HFrEF = heart failure with reduced ejection fraction; HR = heart rate; HRR = heart rate recovery; MVA = multivariable linear regression analysis; $O_2$ = oxygen; OSA = obstructive sleep apnea; PRO = patient-reported outcome; PSQI = Pittsburgh Sleep Quality Index; PTSD = post-traumatic stress disorder; QoL = quality of life; TLC = total lung capacity; SOBQ = University of California San Diego Shortness of Breath Questionnaire; UVA = univariable linear regression analysis

**Table 3B:** Multivariable models for cancer specific QoL (C30 summary score)

|  | Model 1* |  |  | Model 2** |  |  | Model 3† |  |  |
| --- | --- | --- | --- | --- | --- | --- | --- | --- | --- |
| Variable | $\beta$ (95% CI) | Partial R<sup>2</sup> | P value | $\beta$ (95% CI) | Partial R<sup>2</sup> | P value | $\beta$ (95% CI) | Partial R<sup>2</sup> | P value |
| Age, year | N/A | N/A | N/A | 0.03<br>(-0.18, 0.23) | 0.001 | 0.79 | 0.02<br>(-0.20, 0.24) | 0.00 | 0.87 |
| Smoking history | N/A | N/A | N/A | F-statistics | 0.03 | 0.41 | F-statistics | 0.02 | 0.46 |
| Sex (F/M) | N/A | N/A | N/A | N/A | N/A | N/A | -1.43<br>(-9.68, 6.83) | 0.002 | 0.73 |
| White race (N/Y) | N/A | N/A | N/A | N/A | N/A | N/A | 0.26<br>(-5.98, 6.51) | 0.00 | 0.93 |
| BFI, point | -0.36<br>(-0.48, -0.24) | 0.36 | <0.001 | -0.36<br>(-0.48, -0.24) | 0.35 | <0.001 | -0.36<br>(-0.49, -0.24) | 0.35 | <0.001 |
| HADS depression, point | -1.21<br>(-1.79, -0.63) | 0.20 | <0.001 | -1.22<br>(-1.81, -0.64) | 0.21 | <0.001 | -1.21<br>(-1.81, -0.62) | 0.20 | <0.001 |
| PSQI, point | -0.47<br>(-0.92, -0.02) | 0.06 | 0.04 | -0.48<br>(-0.94, -0.02) | 0.06 | 0.04 | -0.49<br>(-0.97, -0.02) | 0.06 | 0.04 |
| SOBQ, point | -0.11<br>(-0.21, -0.01) | 0.06 | 0.04 | -0.11<br>(-0.22, -0.003) | 0.06 | 0.045 | -0.11<br>(-0.22, -0.001) | 0.06 | 0.048 |
| Overall model | R<sup>2</sup> = 0.85, p < 0.001 |  |  | R<sup>2</sup> = 0.85, p < 0.001 |  |  | R<sup>2</sup> = 0.85, p < 0.001 |  |  |

\*Step-wise backwards selection modeling starting with variables with $p < 0.05$ in UVAs: HFREF, OSA, anxiety/depression/PTSD, 6MWD, Post-6MWT BDS, BFI, HADS anxiety, HADS depression, PSQI, SOBQ.
\*\*Including age and smoking history.
†Including age, smoking history, sex, white race.
6MWD = six-minute walk distance; 6MWT = six-minute walk test; BDS = Borg Dyspnea Score; BFI = Brief Fatigue Inventory; C30 = European Organization for Research and Treatment of Cancer QoL Core 30; CI = confidence interval; HADS = Hospital Anxiety and Depression Scale; HFREF = heart failure with reduced ejection fraction; MVA = multivariable linear regression analysis; OSA = obstructive sleep apnea; PSQI = Pittsburg Sleep Quality Index; PTSD = post-traumatic stress disorder; QoL = quality of life; SOBQ = University of California San Diego Shortness of Breath Questionnaire; UVA = univariable linear regression analysis

**Figure 1:**
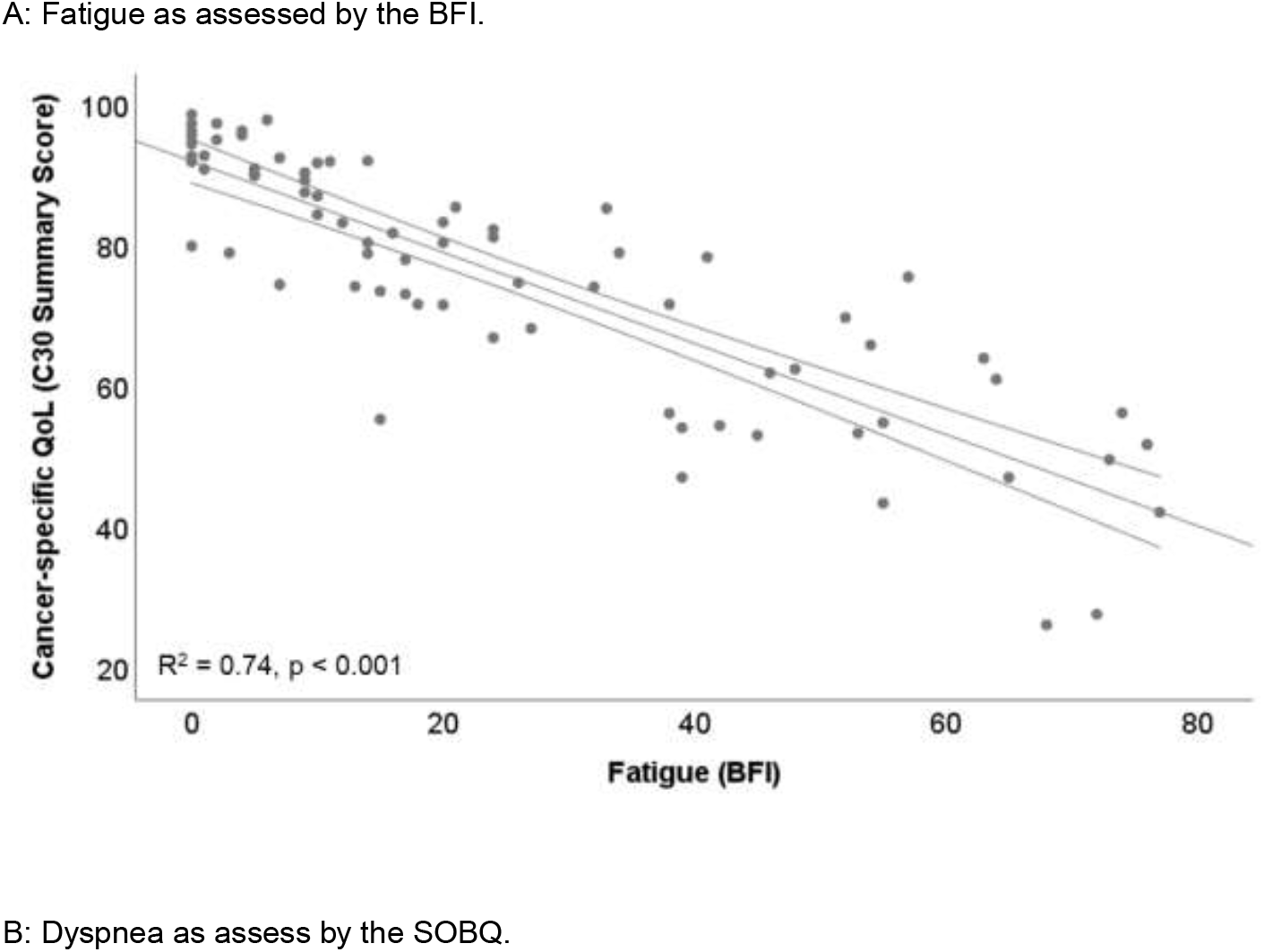
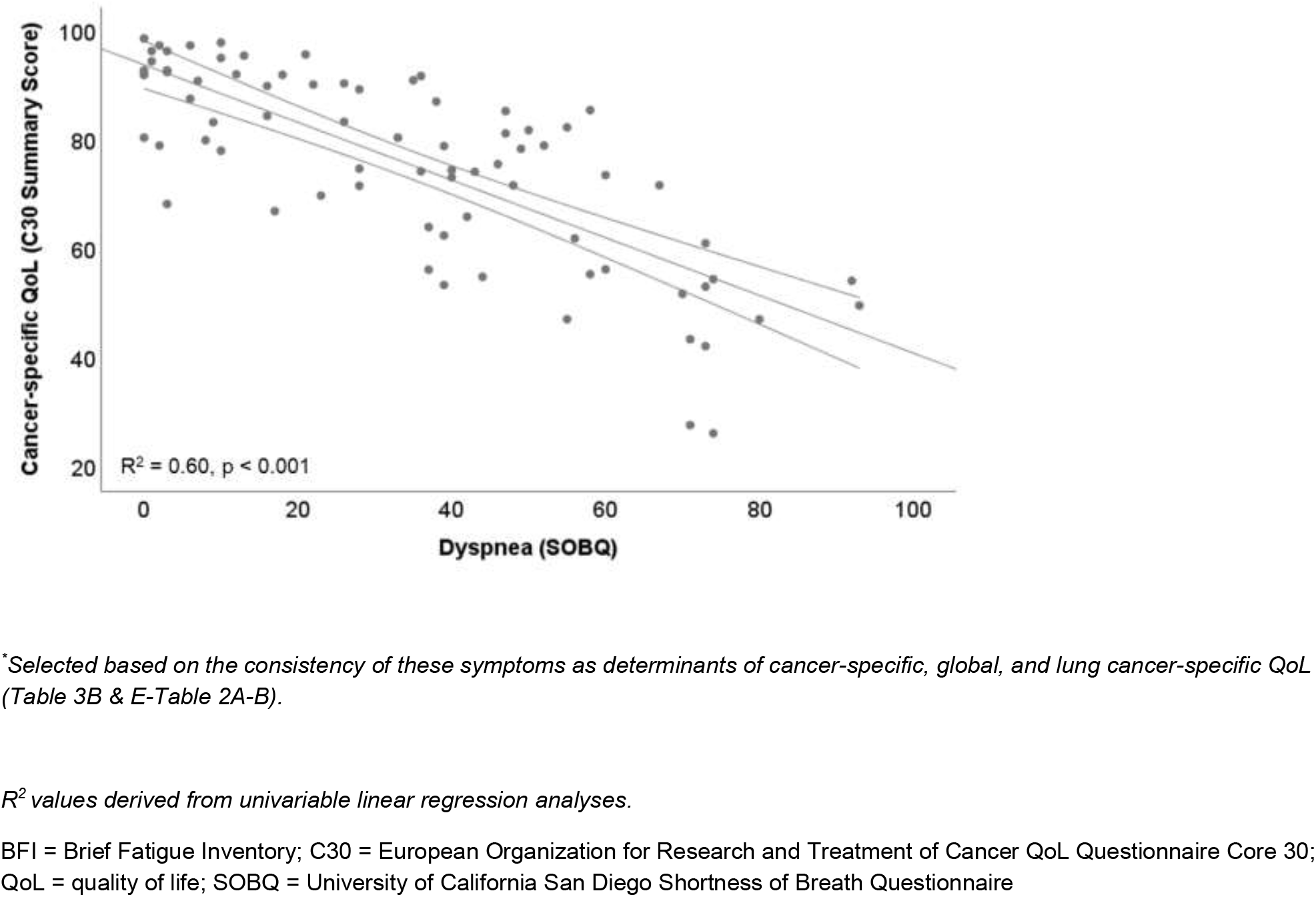
Selected* Determinants of Cancer-specific QoL (C30 Summary Score) A: Fatigue as assessed by the BFI.

## Abbreviations

6MWD: = six-minute walk distance
6MWT: = six-minute walk test;
ATS: = American Thoracic Society;
BDS: = Borg Dyspnea Score;
BFI: = Brief Fatigue Inventory;
BMI: = body-mass index;
CI: = confidence interval;
COPD: = chronic obstructive pulmonary disease;
DL_CO_: = diffusion capacity of the lung for carbon monoxide;
C30/LC13: = European Organization for Research and Treatment of Cancer QoL Questionnaire Core 30/Lung Cancer Module 13;
FEV_1_: = forced expiratory volume in 1 second;
HADS: = Hospital Anxiety and Depression Scale;
HF: = heart failure;
HR: = heart rate;
MVA: = multivariable linear regression analysis;
NSCLC: = non-small cell lung cancer;
O_2_: = oxygen;
PRO: = patient-reported outcome;
PSQI: = Pittsburgh Sleep Quality Index;
QoL: = quality of life;
RTC: = randomized clinical trial;
SD: = standard deviation;
SOBQ: = University California San Diego Shortness of Breath Questionnaire;
US: = United States;
UVA: = univariable linear regression analysis;
VASDHS: = VA San Diego Healthcare System

