## Supplemental Data for "Determinants of Cancer-specific Quality of Life in Veteran Lung Cancer Survivors Eligible for Long-Term Cure"

E-Table 1: UVA – Determinants of Global and Lung Cancer-Specific QoL

|  | C30  Global Health Status/QoL | | | LC13  Summated Score^*^ | | |
| --- | --- | --- | --- | --- | --- | --- |
| Variable | β | R^2^ | P-value | β | R^2^ | P-value |
| Age, year | 0.46 | 0.03 | 0.16 | **-0.14** | **0.06** | **0.04** |
| Sex (F/M) | -20.0 | 0.04 | 0.10 | 0.28 | 0.00 | 0.91 |
| White race (N/Y) | -17.9 | 0.05 | 0.052 | 2.66 | 0.03 | 0.15 |
| BMI, kg/m^2^ | -0.07 | 0.00 | 0.90 | -0.12 | 0.01 | 0.31 |
| Smoking history | F-stats | 0.004 | 0.87 | F-stats | 0.03 | 0.33 |
| Comorbidities (N/Y)  Diabetes mellitus  Atrial fibrillation/flutter  CAD  HFrEF  COPD  OSA  Anxiety/depression/PTSD  Other cancer | -1.70  3.27  -5.14  11.2  -1.11  **14.5**  **18.5**  6.76 | 0.001  0.004  0.01  0.04  0.00  **0.07**  **0.13**  0.02 | 0.78  0.61  0.37  0.08  0.86  **0.02**  **0.001**  0.22 | -0.04  0.45  -0.41  **-3.13**  -0.07  **-3.08**  **-3.34**  0.56 | 0.00  0.002  0.002  **0.08**  0.00  **0.08**  **0.11**  0.003 | 0.97  0.72  0.72  **0.01**  0.96  **0.02**  **0.004**  0.62 |
| Pulmonary function  FEV_1_ % predicted, each  TLC % predicted, each  DL_CO_ % predicted, each | 0.08  -0.04  0.16 | 0.01  0.002  0.03 | 0.45  0.75  0.14 | -0.04  0.02  -0.04 | 0.05  0.005  0.04 | 0.06  0.58  0.07 |
| Lung cancer characteristics  Clinical stage I (N/Y)  Primary treatment  Months since treatment, each | 1.78  F-stats  0.03 | 0.001  0.04  0.002 | 0.79  0.26  0.71 | -1.17  F-stats  0.004 | 0.01  0.07  0.001 | 0.38  0.09  0.76 |
| 6MWT-associated measures  6MWD, each m  Post-6MWT BDS, each point  HR change, each beats/min  O_2_ saturation change, each %  HRR, each beat | **0.05**  **-4.37**  -0.01  0.03  0.21 | **0.07**  **0.24**  0.00  0.00  0.004 | **0.03**  **<0.001**  0.98  0.96  0.60 | -0.01  **0.93**  -0.07  -0.09  0.12 | 0.05  **0.28**  0.03  0.01  0.03 | 0.06  **<0.001**  0.14  0.46  0.15 |
| Symptom-specific PRO, each point  BFI  HADS anxiety  HADS depression  PSQI  SOBQ | **-0.70**  **-2.43**  **-3.27**  **-2.59**  **-0.59** | **0.49**  **0.21**  **0.38**  **0.25**  **0.41** | **<0.001**  **<0.001**  **<0.001**  **<0.001**  **<0.001** | **0.15**  **0.70**  **0.72**  **0.49**  **0.13** | **0.56**  **0.42**  **0.45**  **0.22**  **0.51** | **<0.001**  **<0.001**  **<0.001**  **<0.001**  **<0.001** |

***Bolded*** *variables indicate statistically significant associations at p < 0.05 and selected to enter MVAs.*

*^*^Calculated as a summation of responses from LC13 questions 31 – 42 (maximum score = 48, indicating worst possible QoL).*^1,2^

6MWD = six-minute walk distance; 6MWT = six-minute walk test; BDS = Borg Dyspnea Score; BFI = Brief Fatigue Inventory; CAD = coronary artery disease; COPD = chronic obstructive pulmonary disease; DL_CO_ = diffusion capacity of the lung for carbon monoxide; C30/LC13 = European Organization for Research and Treatment of Cancer QoL Questionnaire Core 30/Lung Cancer Module 13; FEV_1_ = forced expiratory volume in 1 second; HADS = Hospital Anxiety and Depression Scale; HFrEF = heart failure with reduced ejection fraction; HR = heart rate; HRR = heart rate recovery; MVA = multivariable linear regression analysis; O_2_ = oxygen; OSA = obstructive sleep apnea; PRO = patient-reported outcome; PSQI = Pittsburg Sleep Quality Index; PTSD = post-traumatic stress disorder; QoL = quality of life; TLC = total lung capacity; SOBQ = University of California San Diego Shortness of Breath Questionnaire; UVA = univariable linear regression analysis

E-Table 2: Verification of Determinants of QoL

A) MVA – Independent Determinants of Global QoL

|  | Model 1^*^ | | | Model 2*^**^* | | | Model 3*^†^* | | |
| --- | --- | --- | --- | --- | --- | --- | --- | --- | --- |
| Variable | β (95% CI) | Partial R^2^ | P value | β (95% CI) | Partial R^2^ | P value | β (95% CI) | Partial R^2^ | P value |
| BFI, point | -0.41  (-0.66, -0.15) | 0.12 | 0.002 | -0.46  (-0.73, -0.20) | 0.15 | 0.001 | -0.43  (-0.69, -0.17) | 0.14 | 0.002 |
| PSQI, point | -0.97  (-1.94, -0.01) | 0.05 | 0.048 | -1.01  (-1.97, -0.04) | 0.06 | 0.04 | -1.23  (-2.19, -0.27) | 0.09 | 0.01 |
| SOBQ, point | -0.24  (-0.46, -0.02) | 0.06 | 0.03 | -0.22  (-0.44, 0.003) | 0.05 | 0.053 | -0.21  (-0.43, 0.004) | 0.06 | 0.054 |
| Overall model | R^2^ = 0.55, p < 0.001 | | | R^2^ = 0.57, p < 0.001 | | | R^2^ = 0.60, p < 0.001 | | |

*^*^Step-wise backwards selection modeling starting with variables with p < 0.05 in UVAs: OSA, anxiety/depression/PTSD, 6MWD, Post-6MWT BDS, BFI, HADS anxiety, HADS depression, PSQI, SOBQ.*

*^**^Including age (p = 0.73) and smoking history (p = 0.19).*

*^†^Including age (p = 0.73), sex (p = 0.08), white race (p = 0.29), smoking history (p = 0.29).*

B) MVA – Independent Determinants of Lung Cancer-Specific QoL

|  | Model 1^*^ | | | Model 2*^**^* | | | Model 3*^†^* | | |
| --- | --- | --- | --- | --- | --- | --- | --- | --- | --- |
| Variable | β (95% CI) | Partial R^2^ | P value | β (95% CI) | Partial R^2^ | P value | β (95% CI) | Partial R^2^ | P value |
| BFI, point | 0.08  (0.04, 0.12) | 0.16 | <0.001 | 0.08  (0.04, 0.13) | 0.17 | <0.001 | 0.08  (0.03, 0.12) | 0.16 | 0.001 |
| HADS anxiety, point | 0.32  (0.15, 0.50) | 0.16 | 0.001 | 0.34  (0.15, 0.53) | 0.15 | 0.001 | 0.37  (0.17, 0.56) | 0.18 | <0.001 |
| SOBQ, point | 0.05  (0.01, 0.09) | 0.09 | 0.01 | 0.05  (0.01, 0.09) | 0.08 | 0.02 | 0.05  (0.01, 0.09) | 0.08 | 0.02 |
| Overall model | R^2^ = 0.68, p < 0.001 | | | R^2^ = 0.68, p < 0.001 | | | R^2^ = 0.69, p < 0.001 | | |

*^*^Step-wise backwards selection modeling starting with variables with p < 0.05 in UVAs: age, HFrEF, OSA, anxiety/depression/PTSD, post-6MWT BDS, BFI, HADS anxiety, HADS depression, PSQI, SOBQ.*

*^**^Including age (p = 0.70) and smoking history (p = 0.83).*

*^†^Including age (p = 0.89), sex (p = 0.09), white race (p = 0.41), smoking history (p = 0.84).*

6MWD = six-minute walk distance; 6MWT = six-minute walk test; BDS = Borg Dyspnea Score; BFI = Brief Fatigue Inventory; CI = confidence interval; HADS = Hospital Anxiety and Depression Scale; HFrEF = heart failure with reduced ejection fraction; MVA = multivariable linear regression analysis; OSA = obstructive sleep apnea; PSQI = Pittsburg Sleep Quality Index; PTSD = post-traumatic stress disorder; QoL = quality of life; SOBQ = University of California San Diego Shortness of Breath Questionnaire; UVA = univariable linear regression analysis

E-Table 3: UVA – Predictors of Functional Capacity

| Variable | β | R^2^ | P value |
| --- | --- | --- | --- |
| **Age, year** | **-4.58** | **0.10** | **<0.01** |
| White race (N/Y) | 21.3 | 0.003 | 0.67 |
| BMI, kg/m^2^ | -0.25 | 0.00 | 0.93 |
| Smoking status | F-stats | 0.03 | 0.36 |
| Comorbidities (N/Y)  Diabetes mellitus  Atrial fibrillation/flutter  **CAD**  HFrEF  COPD  OSA  Anxiety/depression/PTSD  Other cancer | 7.34  26.9  **68.5**  53.0  41.9  -4.15  10.2  34.4 | 0.001  0.01  **0.07**  0.03  0.02  0.00  0.001  0.02 | 0.82  0.43  **0.02**  0.12  0.21  0.91  0.75  0.24 |
| Pulmonary function  FEV_1_ % predicted, each  TLC % predicted, each  **DL_CO_ % predicted, each**  **Combined vent. abnormalities*^*^*** | 0.96  -0.08  **2.07**  **F-stats** | 0.04  0.00  **0.18**  **0.20** | 0.097  0.92  **<0.001**  **<0.01** |
| Lung cancer characteristics  Clinical stage I (N/Y)  **Primary treatment**  Months since treatment, each | -12.0  **F-stats**  -0.02 | 0.002  **0.18**  0.00 | 0.73  **<0.01**  0.95 |
| 6MWT-associated measures  **HR change, each beats/min**  O_2_ saturation change, each %  Post-6MWT BDS, each point  **HRR, each beat** | **3.54**  1.15  -8.78  **-6.27** | **0.12**  0.002  0.04  **0.12** | **<0.01**  0.71  0.11  **<0.01** |
| Symptom-specific PRO, each point  **BFI**  HADS anxiety  HADS depression  PSQI  **SOBQ** | **-1.54**  1.94  -6.26  -4.41  **-1.85** | **0.08**  0.01  0.05  0.03  **0.15** | **0.01**  0.58  0.057  0.17  **<0.01** |
| EORTC-QLQ-LC13, each point  Dyspnea  Coughing  Hemoptysis  Sore mouth  Dysphagia  Peripheral neuropathy  Alopecia  Pain in chest  Pain in arm or shoulder  **Pain in other parts** | -1.49  0.39  -0.44  -0.70  -1.04  0.06  1.02  0.38  -0.48  **-0.95** | 0.10  0.01  0.004  0.004  0.03  0.00  0.01  0.01  0.01  **0.07** | <0.01  0.48  0.61  0.59  0.14  0.90  0.34  0.51  0.32  **0.03** |
| Quality of Life  **Cancer-specific QoL**  **Global QoL**  Lung cancer-specific QoL | **2.32**  **1.36**  -5.71 | **0.11**  **0.07**  0.05 | **<0.01**  **0.03**  0.06 |

*^*^Defined as a summation of obstructive (FEV_1_/FVC < 70%), hyperinflated (TLC % predicted > 120), and DL_CO_ limitation (DL_CO_ % predicted < 80); range 0-3.*

***Bolded*** *variables indicate statistically significant associations at p < 0.05 and selected to enter MVAs; EORTC-QLQ-LC13 dyspnea not bolded due to redundancy with SOBQ.*

6MWT = six-minute walk test; BDS = Borg Dyspnea Score; BFI = Brief Fatigue Inventory; BMI = body-mass index; CAD = coronary artery disease; COPD = chronic obstructive pulmonary disease; DL_CO_ = diffusion capacity of the lung for carbon monoxide; EORTC-QLQ-LC13 = European Organization for Research and Treatment of Cancer QoL Questionnaire Lung Cancer Module 13; FEV_1_ = forced expiratory volume in 1 second; HADS = Hospital Anxiety and Depression Scale; HFrEF = heart failure with reduced ejection fraction; HR = heart rate; HRR = heart rate recovery; MVA = multivariable linear regression analysis; O_2_ = oxygen; OSA = obstructive sleep apnea; PSQI = Pittsburg Sleep Quality Index; PTSD = post-traumatic stress disorder; QoL = quality of life; TLC = total lung capacity; SOBQ = University of California San Diego Shortness of Breath Questionnaire; UVA = univariable linear regression analysis
